# Efflux mediated chlorpyrifos tolerance in *Escherichia coli* BL21(DE3)

**DOI:** 10.1101/2020.10.08.330753

**Authors:** Aswathi Aswathi, Ashok Pandey, Rajeev K Sukumaran

## Abstract

Bacteria are continually challenged with variety of synthetic chemicals/xenobiotics in their immediate surroundings, including pesticides. Chlorpyrifos is one of the most commonly used organophosphate pesticides in the world. The non-environmental strain of *Escherichia coli,* BL21 (DE3) displayed high tolerance to chlorpyrifos but with no/negligible degradation. The intrinsic resistance mechanisms that aid the organism in its high tolerance are probed. Efflux pumps being ubiquitous in nature and capable of conferring resistance against wide variety of xenobiotics were found to be over-expressed in the presence of CP. Also, an efflux pump inhibitor PAβN increased the susceptibility of *E. coli* to chlorpyrifos due to the intracellular accumulation of CP. The tripartite efflux pump EmrAB-TolC with increased expression in both transcript and protein on CP exposure, might play a major role in CP tolerance. The transcriptional regulators involved in multidrug resistance along with transporters belonging to all the major families conferring antimicrobial resistance were up-regulated. Also up-regulated were the genes involved in phopshonate metabolism and all the genes in the copper or silver export system. The common resistance mechanisms i.e, activation of efflux pumps between CP, antibacterial metals and antibiotics resistance might result in cross-resistance, ultimately increasing the prevalence of multidrug resistant strains, making infections hard to treat.

## 1. Introduction

Bacteria evolve various complex systems for their survival in presence of toxic compounds. One such system involving efflux pumps confer multidrug resistance (mdr), making infections hard to treat (Tanabe et al., 2009). Efflux pumps are ubiquitous in nature and perform one to many functions such as conferring resistance to antimicrobial drugs, and are required for virulence and biofilm formation. They can be broadly classified into two groups, namely primary and secondary transporters. Those that depend on ATP for energy are the primary transporters such as ATP-binding cassette (ABC) transporters and those that use the membrane electro-potential such as major facilitator superfamily (MFS), small multidrug resistance (SMR) family, resistance nodulation division (RND) family and multidrug and toxic compound extrusion (MATE) family, are the secondary transporters (Paulsen, 2003; Blair et al., 2014).

The tripartite efflux pumps capable of transporting toxic compounds across both the outer and inner membranes also play a vital role in multi-drug resistance (mdr). Tripartite efflux system consists of an inner membrane transporter belonging to RND or MFS families that couples the energy of proton gradient to haul molecules across membranes and an outer membrane channel protein (OMP) along with a membrane fusion protein (MFP) that binds the OM and IM domains together in periplasm (Tanabe et al., 2009). Of particular interest is the outer membrane channel protein TolC, required for the functioning of all RND drug transporters, namely AcrAB, AcrD, AcrEF, MdtEF and MdtABC (Elkins and Nikaido, 2002; Kobayashi et al., 2001; Nishino and Yamaguchi, 2001) along with two MFS transporters (emrAB, emrKY) and an ABC transporter MacAB (Kobayashi et al., 2001; Lomovskaya and Lewis, 1992; Pasqua et al., 2019), involved in xenobiotic resistance in *E. coli*. Some drug transporters from MFS, SMR and MATE families, namely emrE, mdfA and mdtK are TolC-independent transporters (Nishino et al., 2003).

The active transport of chemicals by the efflux pumps is characterised by reduced intrinsic susceptibility, possible cross resistance to chemically diverse classes and augmenting enhanced resistance adaptations. Thus, inhibition of bacterial efflux pumps by efflux pump inhibitors is a promising strategy to increase the accumulation of pump substrates to a lethal concentration and also to diminish the acquired/adaptive resistance (Mahamoud et al., 2007). Phenylalanine-arginine β-naphthylamide (PAβN) is a broad spectrum peptidomimetic efflux pump inhibitor mostly effective against resistance to fluoroquinolone antibiotics in *P. aeruginosa*. It is a substrate of efflux pumps, functioning as a competitive inhibitor of pump function and can be used to study the substrates of efflux pumps (Lamers et al., 2013; Lomovskaya et al., 2001; Sáenz et al., 2004).

Bacterial responses to xenobiotics acting as antimicrobials have been widely studied, the most prevalent being antibiotics and non-antibiotics like pesticides, being the least. The ability to survive under difficult environmental conditions combined with the extensive knowledge on its genetics, transcriptome, proteome and metabolome makes *Escherichia coli* a well-established model organism for bacterial molecular studies (Idalia and Bernardo, 2017). The transcriptomic studies on the stress responses of *E coli* to pesticides such as glyphosate (Lu et al., 2013), paraquat (Allen and Griffiths, 2012) and *p-*nitrophenol (Chakka et al., 2015) were recently reported. Though CP is a widely used and moderately toxic insecticide with several reports on its bacterial degradation, the existing literature on the molecular mechanism aiding bacterial response to chlorpyrifos (CP) are scarce to none.

We studied the changes in gene expression that accompanied the exposure of a non-environmental strain of *E. coli* BL21(DE3) to the organophosphate (OP) pesticide-chlorpyrifos (CP), to elucidate the inherent mechanisms that assisted in its high tolerance against the pesticide.

## 2. Materials and Methods

### 2.1. Chemicals and media

The *E. coli* was grown in mineral salt media (MSM, pH 7.0) containing (g/L) KH_2_PO_4_- 1.0; K_2_HPO_4_-2.0; MgSO_4_.7H_2_O-1.0; (NH_4_)_2_SO_4_-5.0; yeast extract-0.5 and glucose 1.0. Technical grade CP (96%) was obtained from the Indian Institute of Chemical Technology, CSIR, Hyderabad, India and the broad spectrum efflux pump inhibitor - Phenylalanine-arginine β-naphthylamide (PAβN) was purchased from Sigma Aldrich, India. A chlorpyrifos stock of 500 g/L in methanol (HPLC grade) and PAβN stock of 20 g/L in sterile water was used until otherwise mentioned and subsequently added to the medium to obtain desired concentrations.

### 2.2 Determining the effect of chlorpyrifos on bacterial growth

*E. coli* strain BL21 (DE3) and *Pseudomonas nitroreducens* strain AR-3, a chlorpyrifos degrader, were evaluated for their growth in mineral salt medium (MSM), supplemented with 10, 25, 50 or 100 mg/L of chlorpyrifos. A 2% (v/v) of an overnight grown bacterial culture was used as inoculum. Growth was measured as optical density at 600 nm (OD_600_) and monitored every 2h till the 12^th^ hour. Growth in CP containing media was compared against controls without any CP addition. Mean values of triplicates are reported

### 2.3. Effect of efflux pump inhibition on bacterial growth in presence of chlorpyrifos

Overnight grown culture of *E. coli* BL21(DE3) in Luria Bertani medium (25ml in 100ml Erlenmeyer flask) was washed three times and suspended in 25ml of sterile N-saline (0.9% NaCl). The suspension was used at 1% (v/v) level for inoculating 25 mL sterile MSM taken in 100 ml Erlenmeyer flasks. Chlorpyrifos (100 mg/L) was added from the stock solution, followed by efflux pump inhibitor PAβN. The growth was monitored as OD_600_ in a UV spectrophotometer (UV-1800, Shimadzu).

### 2.4. Chlorpyrifos accumulation in *E. coli* cells in presence of efflux pump inhibitor

Chlorpyrifos accumulation in *E coli* cells, in the presence of efflux pump inhibitor was studied in 250 mL Erlenmeyer flasks containing 50 mL of MSM supplemented with 1000 mg/L CP, and either 40 mg/L or no PAβN. Flasks were inoculated with 1% (v/v) inoculum prepared as in section 5.2.2, and were incubated at 37°C for 12h in a rotary shaker incubator at 180 rpm. Cultures were extracted with an equal volume of olive oil to remove CP and the aqueous layer containing pesticide-free cells were pelleted (3300×g for 8 min, 4°C) and washed 6× with sterile N-saline. The cell free supernatant was lyophilized and stored at 4°C as control. The cell pellet was re-suspended in N-saline, sonicated on ice and the resultant lysate was dehydrated to a powder by lyophilisation. One millilitre methanol was added to the total powdered lysate, thoroughly mixed for 30min, followed by centrifugation at 3300×g for 10 min at 25 °C. The supernatant was recovered and evaporated to dryness in a vacuum concentrator; and the residues were re-dissolved in 1.0 mL acetonitrile. This was used to determine the cellular CP accumulation by HPLC analysis (Aswathi et al., 2019).

### 2.5. Determining the differentially expressed genes in *E. coli* in response to chlorpyrifos by transcriptome analysis

Genes differentially expressed in *E. coli* in response to CP was determined through a comparative transcriptome profiling of CP treated and untreated bacterial cultures. Twenty-five millilitres of an actively growing (4h old, OD_600_ ~ 0.6) LB culture of *E. coli* was washed three times (3300×g for 8 min, 4°C) with equal volume of sterile N-saline and suspended in 25ml of fresh mineral media. The cell suspension (2%v/v) so prepared was used for inoculating 50ml MSM in 250ml flasks supplemented with 500 mg/L of CP and was grown for 3 h at 37 °C and 180 rpm on a rotary shaker incubator until mid-exponential phase. The control consisted of cells grown similarly but without addition of CP in the medium. The culture broth (4.0 mL) after 3h of incubation was pelleted (3300 x g for 8 min, 4°C) and RNA was isolated using Trizol® Plus RNA Purification Kit (Ambion, Life Technologies, USA). The quality and quantity of RNA was determined by agarose gel electrophoresis and also using a NanoDrop ND-2000 spectrophotometer. The library construction and RNA sequencing were outsourced (OmicsGen LifeSciences Pvt Ltd, Kochi, India); carried out using the Illumina HiSeq 2000 platform.

Quality parameters for sequenced reads were determined using FastQC (Cock et al., 2009). The resultant reads were aligned to the reference genome - *Escherichia coli* Str.K12 (NC_000913.3) using HISAT2 (Pertea et al., 2016). The individual transcripts from RNA-seq reads, aligned to the genome were assembled by StringTie (Pertea et al., 2015). The assembly was then screened for transcripts/genes that were differentially expressed using Cuffdiff v2.2.1 (Trapnell et al, 2010) that calculated the FPKM (Fragments Per Kilo bases per Million reads) using the following formula

FPKM = Total exon fragments / [Mapped reads (Millions) * exon length (kb)] (Trapnell et al., 2012).

The differentially expressed genes/transcripts retrieved were annotated using UniProt (UniProt Consortium, 2018), KEGG (Kanehisa et al., 2010) and Gene Ontology (GO) databases (Camon et al., 2004). The significant (p<0.05, log FPKM >1) differentially expressed genes were further mapped to pathways using KEGG/EcoCyc pathway analysis databases.

### 2.6. Comparative proteome profiling to determine differential protein expression in response to chlorpyrifos

Culturing of the bacterial cells in presence of CP and control cultures were performed as described in section 5.2.5. The mid exponential phase (3h) culture broth (50ml) was extracted with equal volume of olive oil to remove residual CP, and the aqueous layer containing bacterial cells was pelleted (3300×g for 8 min, 4°C), washed six times with phosphate buffered saline (PBS) and resuspended in 0.6ml of PBS supplemented with protease inhibitory cocktail (Roche) as per manufacturer’s specification. The suspension was sonicated on ice, and the resultant lysate was centrifuged at 13,400×g, at 4°C for 20 min. The supernatant was estimated for protein by Bradfford assay, and 100 μg of protein after trypsin digestion was analysed by liquid chromatography tandem mass spectrometry - LC MS/MS (Vineetha et al., 2020).

## 3. Results and Discussion

### 3.1. Efflux pumps inhibition leads to growth reduction in presence of chlorpyrifos

Efflux pump inhibition with 40 mg/L inhibitor (PAβN) resulted in cell (*E. coli*) growth being negatively affected in presence of chlorpyrifos (Fig 5.1). The efflux pump inhibitor (EPI) when supplemented alone in the medium, resulted in considerable growth reduction compared to the un-treated control, indicating efflux pump inhibition can cause growth reduction in the bacteria even in the absence of CP. PAβN treatment was reported to compromise the outer membrane barrier resulting in release of bulky folded periplasmic proteins to the outside whose effects partially reversed by Mg^++^. A significant but non-lethal disruption of the membrane potential due to depolarization of inner membrane was also observed upon the treatment with PAβN (Lamers et al., 2013). These might explain the reduced growth of cells on efflux pump inhibitor (PAβN) alone.

**Figure 5.1:**
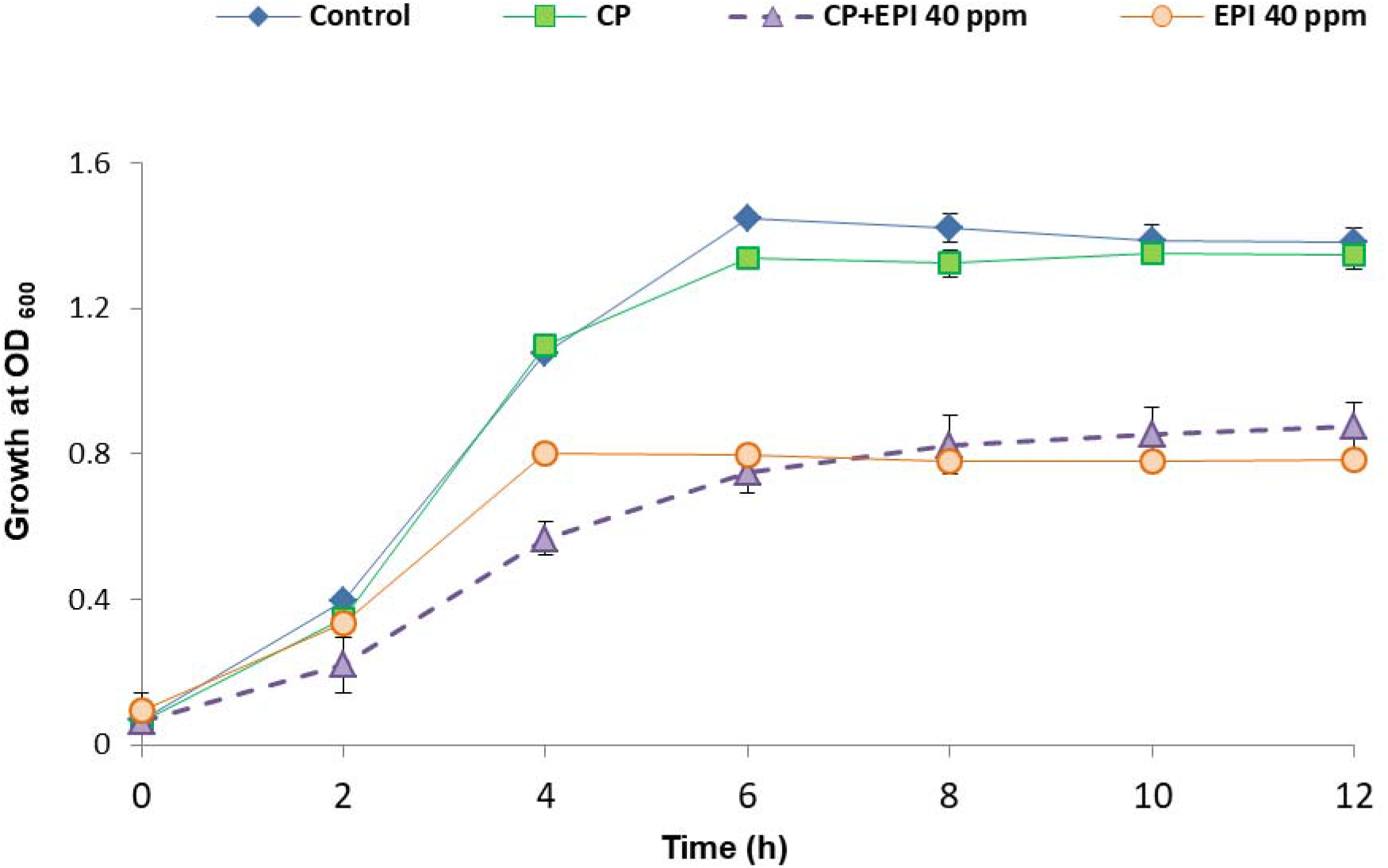
The effect of efflux pump inhibition on the growth of CP stressed *E. coli.* Growth measured as optical density (OD_600_). Control group does not have EPI or CP, CP and EPI were supplemented at 100mg/L and 40mg/L respectively. Values are means of three replicates.

Nevertheless, when the organism was challenged with chlorpyrifos in presence of the EPI, there was a much reduction in the growth of the bacterium, taking double the time (8h instead of 4h) to attain the maximum cell density (OD_600_), against those supplemented with EPI (PaβN) alone. In both the cases (EPI or EPI +CP), the maximal cell densities achieved were much lesser than the control. While CP alone at 100 mg/L did not cause any significant growth inhibition, when it was given in presence of 40mg/L EPI, the growth was seriously affected. The observation that CP does not affect the bacterial growth while the efflux pumps are active; indicates the involvement of efflux pumps in CP tolerance of *E. coli* and preferentially mediated by disallowing intracellular CP concentration from reaching harmful levels by pumping it out. Also, the efflux mechanisms could be a generic line of defence against xenobiotics by the bacterium; since *E. coli* primed with non-antibiotics like salicylate, which is known to upregulate genes for efflux pumps (Cohen et al, 1993) is reported to improve resistance to antibiotics (Laudy et al, 2016). If the CP tolerance would have been mediated through efflux of the compound, it could be expected to accumulate in the cells when the efflux pumps are blocked using the inhibitor. Further studies were undertaken to monitor the accumulation of CP inside the cell upon blocking the efflux pump using PAβN.

### 3.2 Efflux pump inhibition resulted in cellular chlorpyrifos accumulation

Cellular accumulation of chlorpyrifos was monitored in presence and absence of EPI (PAβN). A control was run in parallel without EPI and CP. Maximum detectable cellular chlorpyrifos accumulation (~7 mg/L) was observed when the bacteria were exposed to the pesticide in presence of the broad spectrum efflux pump inhibitor PAβN. Since the amount of biomass (wet weight) was quite low in culture treated with CP+EPI compared to that of CP treatment alone, the biomass amount in the two sets of experiments were normalized for monitoring CP accumulation. All biomass was initially extracted with olive oil followed by washes in saline to remove any residual CP; which was ensured by measuring CP levels of the final wash liquid.

Efflux pump inhibition is known to increase the accumulation of its substrate in the cells. While supplementing CP in culture medium did result in its presence inside the cells, the intracellular CP concentration (measured as peak area) was 45 fold higher in EPI containing medium compared to that without it (Fig 5.2). This indicates the possibility that CP might act as a substrate of efflux pumps, and the organism’s tolerance to CP could be mediated through its removal from the cells by efflux pumps. It was shown that the EPIs reduced the activity of MDR pumps which are mostly associated with antimicrobial resistance, thereby impeding the emergence of antibiotic resistance that occurred as a result of mutation in MDR regulator elements, or by reducing the intrinsic resistance (Vila and Martinez, 2008). PAβN strongly inhibits efflux activity and acts mainly as efflux pump inhibitor at low concentrations. At high concentrations, it not only inhibits efflux but also destabilizes the outer membrane and influence permeability (Misra et al., 2015).

**Figure 5.2:**
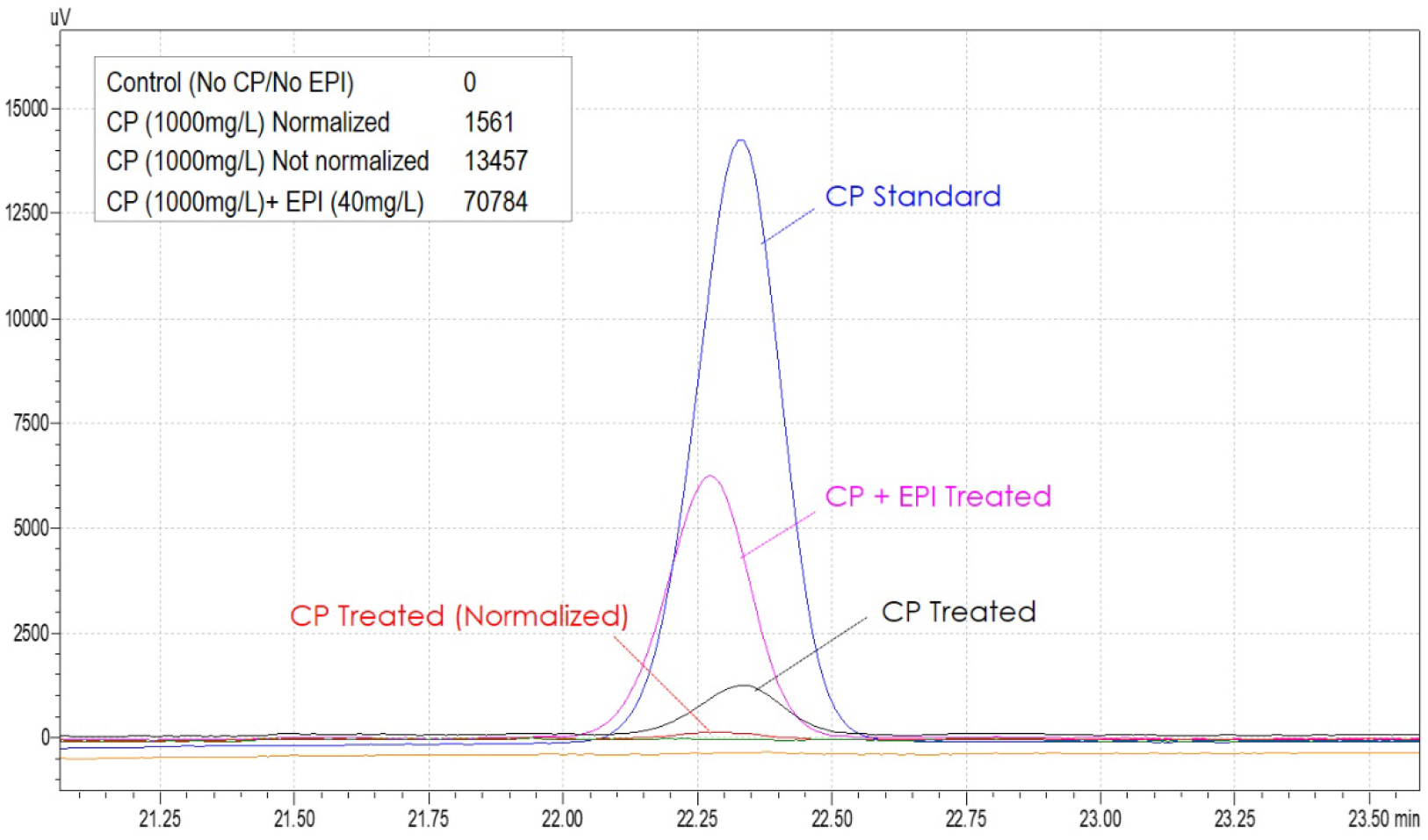
Cellular accumulation of chlorpyrifos in *E. coli* in the presence of efflux pump inhibitor (PAβN). CP levels expressed as area under peak is shown as inset.

### 3.3 Elucidation of the chlorpyrifos tolerance mechanism of *E. coli* by profiling of differentially expressed genes (DEGs) and proteins

The *E. coli* strain BL21(DE3) tolerated high concentrations of CP, but at the same time there was no evidence for degradation of the pesticide by the organism. The studies performed on efflux pumps above, proved that CP is indeed cleared from the cells through an efflux mechanism, and disrupting the efflux pump function makes it susceptible to CP, as can be deduced from its influence on bacterial growth. Therefore, it was apparent that tolerance to the pesticide is a complex phenomenon and possibly involving changes in gene/protein expression. The organism was therefore studied for changes in gene expression induced by the pesticide, performed through RNA-Seq. A total of 3857 differentially expressed genes (DEG) were identified, of which 87% showed no significant differences in expression between CP treated and untreated (control) while 5.8% were significantly up-regulated (≥1 log_2_ fold change) and 6.9% of the genes were significantly down-regulated (≤ −1 log_2_ fold change) as shown in Figure 5.3. Upon pathway analysis by KEGG, 57 pathways were identified as up-regulated with an increase in expression of transporters, ribosomal and quorum sensing genes and noncoding RNAs. The key pathways up-regulated were those involved in signalling and cellular processes, metabolism and environmental information processing.

**Figure 5.3:**
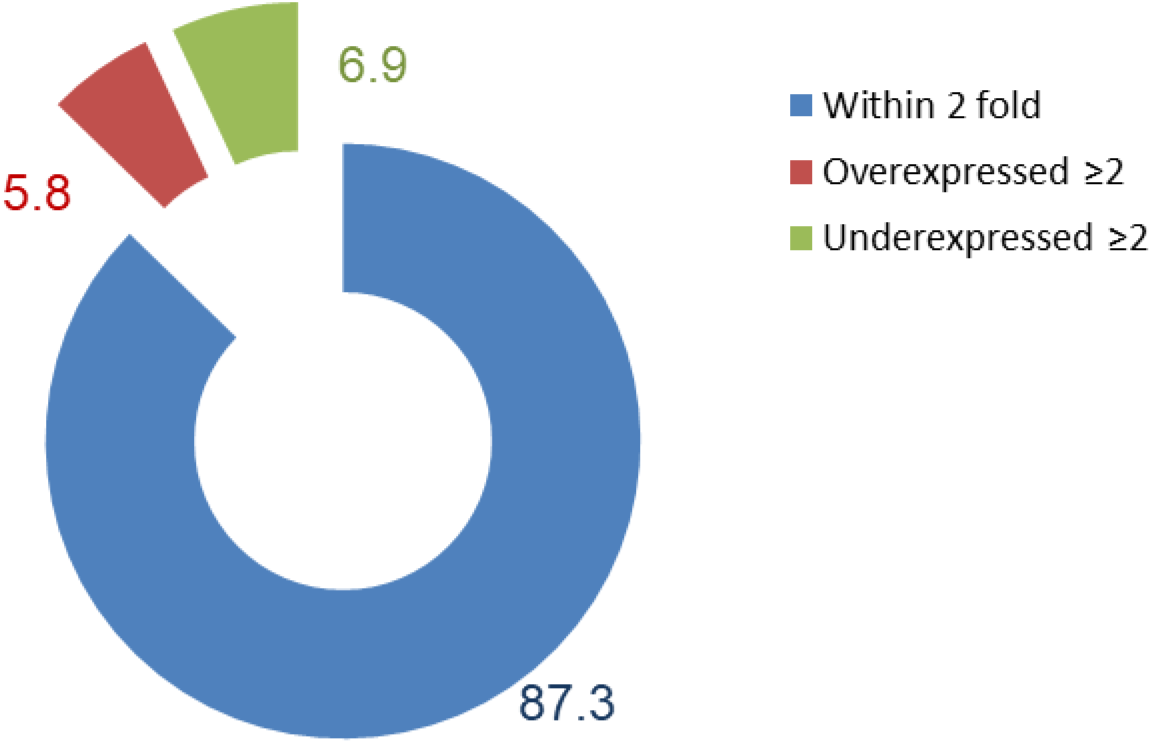
Percentage of DEGs on treatment with 500 mg/L of CP against an untreated control.

#### 3.3.1. *E. coli* exposed to chlorpyrifos shows increased expression of efflux pumps

Drug transporters working in a coordinated manner forms the first line of defence against the toxins such as antibiotics by preventing their cellular concentration from reaching the lethal range until adaptations initiate to aid survival. Gene expression analysis of CP stressed *E. coli* revealed that two major types transporters were over-expressed, which were the single component transporters that move compounds from cytoplasm to periplasm and the multi component transporters-mainly the TolC-dependent transporters, capable of transporting chemicals across both the inner and outer membranes. These transporters are recognized to provide intrinsic resistance to bacteria against xenobiotics (Schuldiner, 2018). The major single component pumps differentially expressed in response to chlorpyrifos were those belonging to two families SMR (Mdt JI) and MATE pump Mdt K. The TolC-dependent transporters showing more than 2-fold differential expression included Emr AB, MdtE and EmrK (Figure 5.4). EmrAB-TolC might play a major role in tolerance of *E. coli* to the pesticide chlorpyrifos as all the constituent components of the tripartite pump were significantly over-expressed.

**Figure 5.4:**
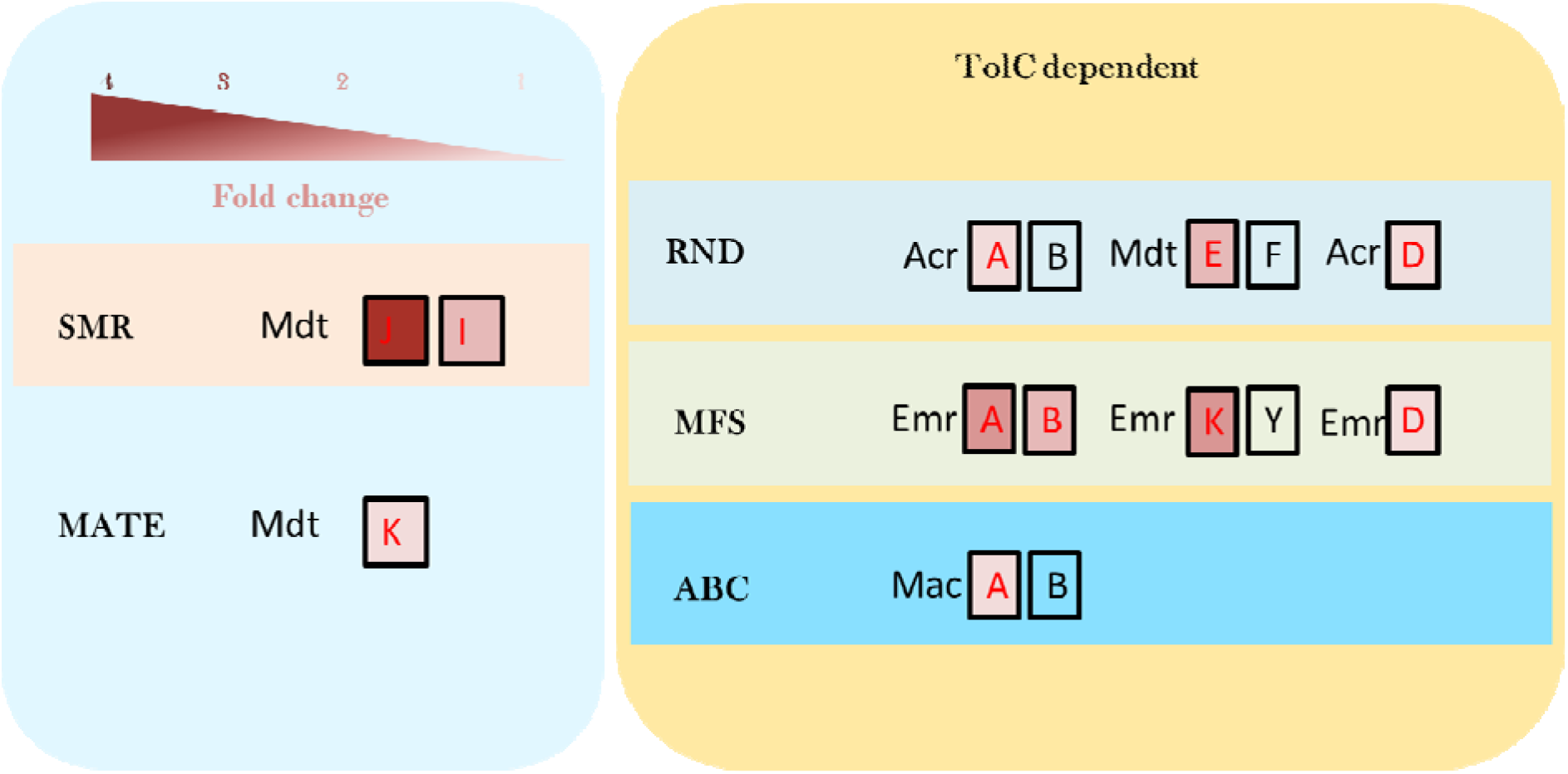
Efflux pumps/transporters overexpressed in *Escherichia coli* on exposure to CP. The efflux pumps belonging to the superfamilies RND, MFS and ABC transporters are dependent on a membrane channel protein TolC for their function while SMR and MATE transporters are TolC-independent. The fold change is denoted by a gradient arrow(red).

#### 3.3.2. Effect of chlorpyrifos on the expression of major transporters and their regulators involved in the efflux of drugs/xenobiotics in *E. coli*

Chlorpyrifos exposure enhanced the expression of transporters and their regulators majorly involved in the transport of metal ions, phosphonate import and xenobiotic transport. Bacterial transporters/efflux pumps belong to five known super families namely- resistance nodulation division (RND), major facilitator super family (MFS), ATP binding cassette (ABC) transporters, small multidrug resistance (SMR) and multi-antimicrobial extrusion (MATE) transporters. MFS and RND pumps are the most abundant with relatively narrow substrate ranges for the former and wide spectrum of substrate ranges for the latter. Multi drug resistance (MDR) transporters are transporters capable of translocating more than three classes of antibiotics and their expression is regulated by the MDR regulon (Blair et al., 2014). Exposure to the organophosphate pesticide-CP enhances the expression of transporters belonging to all the five known transporter super families, with MFS transporters showing the maximum numbers (14), followed by RND (3). The significantly overexpressed transporters from MFS superfamily included emrA, emrB, emrK, yebQ, fsr, ydhC, mdtL, yajR and ydiM along with the RND transporters mdtE and cusA, and SMR transporters mdtJ and mdtI. Along with the transporters, there was a significant upregulation of porins (ompX and ompA) and outer membrane channel protein tolC. The multi-drug resistance (MDR) regulators marA, marB, marR and mprA were significantly up-regulated. The genes phnD, phnI, phnG and phnJ from the phosphonate (Phn) operon were also upregulated (Figure 6.5). The MDR regulon controls the expression of MDR transporters involved in antimicrobial transport and hence culminating in antimicrobial resistance (AMR). The genes involved in transport which have significant differences (≥ 1.5 fold) in expression on CP exposure are shown in Figure 6.6.

It was observed that the entire operon involved in copper/silver resistance (cus system) was over-expressed on CP stress. This included the membrane component of copper efflux pump cusA, membrane fusion protein cusB, outer membrane component cusC, periplasmic copper and silver binding protein (cusF) and the two-component regulatory system cusR/cusS. The copper transporter copA, multicopper oxidase cueO and the cytochrome bo (3) ubiquinol terminal oxidase cyoA involved in copper tolerance were also significantly up-regulated (Table 5.1). The cusCBA system transports Ag(I) and Cu(I) across the membranes to protect the bacterial cells from potential toxicity caused by the accumulation of free metal ions prevalent during the stress (Mealman et al., 2012). Improvement in the osmotolerance of *E. coli* on activation of copper efflux genes was suggested by Xiao et al., (2017). Pal *et al*. (2017) suggested by reviewing antimicrobial metal resistance that antibacterial metals and biocides also contribute to increasing antibiotic resistance possibly when genes for both are collocated or when the resistance mechanisms are similar, like in the case of efflux pumps, which ultimately lead to cross resistance. Since the transcriptome profiling together with EPI experiments suggests the efflux mediated CP tolerance of *E. coli*, the possible cross-resistance against antimicrobial metals and antibiotics cannot be ruled out.

**Table 5.1:**
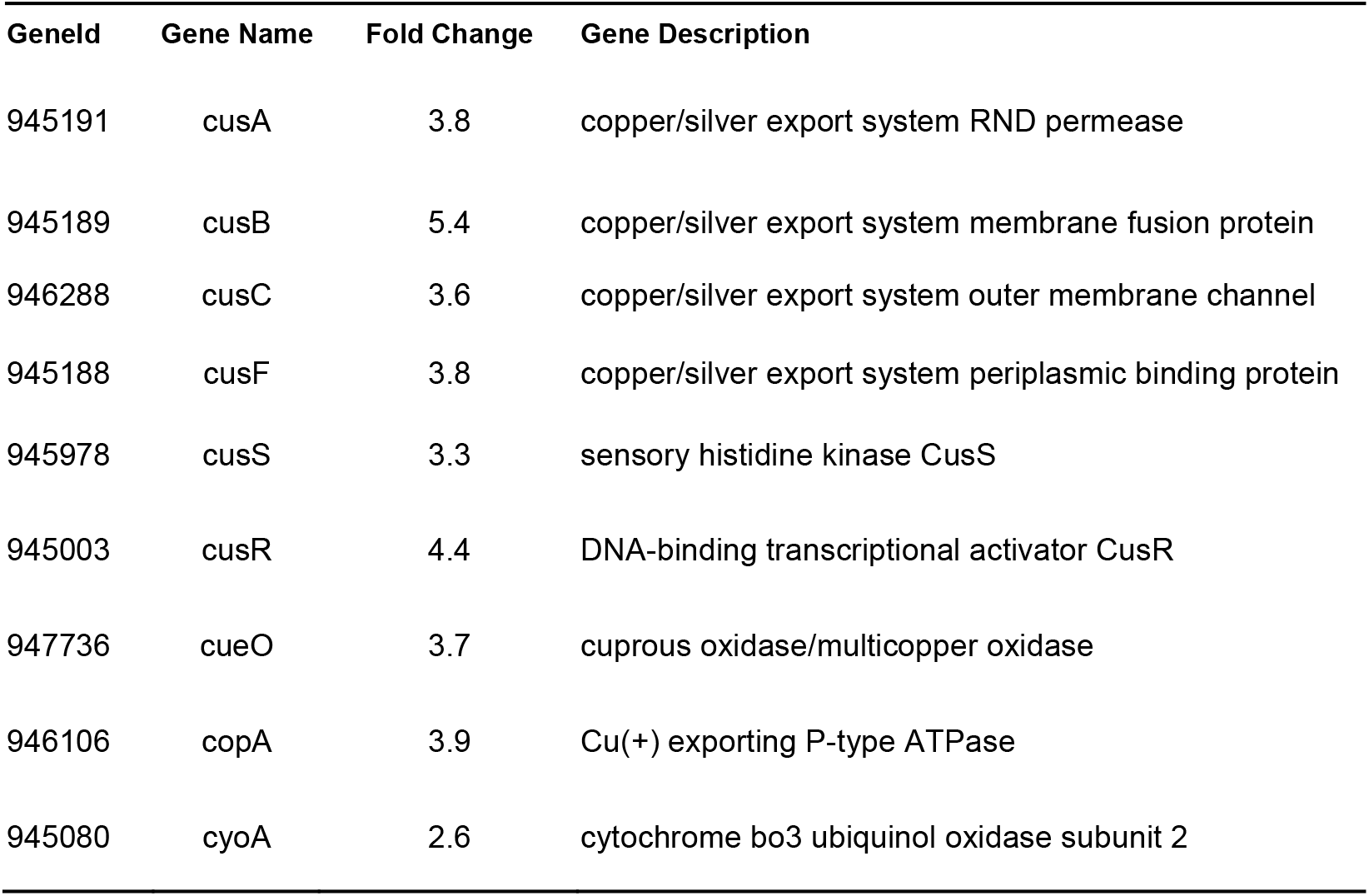
The copper/silver resistance proteins over-expressed during CP stress

CP stress also significantly induced the phosphonate (Pn) transport system, the component genes of which are located in the phn operon. The Pn transport system is involved in the uptake of Pn (P valence +3) which are organic phosphates (Stasi et al., 2019). The genes upregulated included phnD (3.9-fold); the periplasmic binding protein of an ABC type Pn transporter, and the genes that coded for the enzymes involved in Pn metabolism - phnI, phnG and phnJ. This is particularly interesting as the gene phnI is known to code for the enzyme that catalyses the formation of the intermediate ribose-1-phosphonate-5-triphosphate (RPnTP) through displacement of adenine from ATP in presence of PhnG, PhnH and PhnL. A pyrophosphate is cleaved from RPnTP to form ribose-1-phosphonate-5-phosphate (PRPn) by PhnM. The enzyme that cleaves the C-P bond in PRPn to form ribose-1,2-cyclic-phosphate-5-phosphate (PRcP) and methane is the critical reaction and is catalysed by PhnJ (Kamat et al, 2011). Incidentally this gene is also upregulated with chlorpyrifos stress. Previous reports had suggested that phnCDE is also capable of transporting Pi esters and Pi (Metcalf and Wanner, 1991; Stasi et al., 2019). While C-P lyase complexes can have substrate specificities towards different phosphonates (Sviridov et al, 2012), and the Phn operon has as yet undescribed functionalities, it is unclear whether CP can be processed through the phosphonate pathway in *E coli*. While it is rather speculative at this point, the upregulation of Phn operon may be consequence of an increase need for phosphorous resulting from its reduction in cells due to the changes facilitating CP efflux.

#### 3.3.3. CP treatment induces the expression of multidrug resistance (MDR) regulon and transporters

Over-expression of the efflux pumps has been regarded as an intrinsic defence mechanism in Gram-negative bacteria that culminates in increased antibiotic resistance. Those efflux pumps capable of transporting more than one structural class of antimicrobial drugs/antibiotics are called Multi Resistance Drug Resistance (MDR) transporters. They play a major role in antibiotic resistance (Blair et al., 2014). The bacterial resistance to xenobiotics or antimicrobials involve diverse mechanisms whereby bacteria try to minimise the toxic effects by altering the membrane permeability with the help of porins or expel it by the activation of efflux pumps. Once the xenobiotic reaches a tractable intracellular concentration, the other detoxification mechanisms such as enzymatic degradation/transformation to a less toxic form gets into action.

We observed the up-regulation of three porins, namely ompA, ompG and ompX and MDR transporters in *E. coli* in response to CP stress (Figure 5.5). The multiple antibiotic resistance (mar) operon of *E.coli* modulates the expression of efflux pumps and porins *via* two transcription factors marA and marR (Ariza et al., 1994; Sharma et al., 2017). The genes marA, marB and marR belonging to the mar operon were upregulated in the *E.coli* exposed to CP. A negative regulator of the multidrug operon emrAB (mprA) was also significantly up-regulated. Analysis of the antimicrobial resistance genes/mechanisms from the transcriptome performed using comprehensive antibiotic resistance database (CARD) revealed the activation of three major antimicrobial resistance mechanisms in CP stressed *E.coli*, namely antibiotic efflux (emrA, emrB, emrR, marA, emrK, mdtE), reduced permeability to antibiotics (marA, ompA) and antibiotic target alteration (marR). The genes attributed for fluoroquinolone resistance were the most abundant.

**Figure 5.5:**
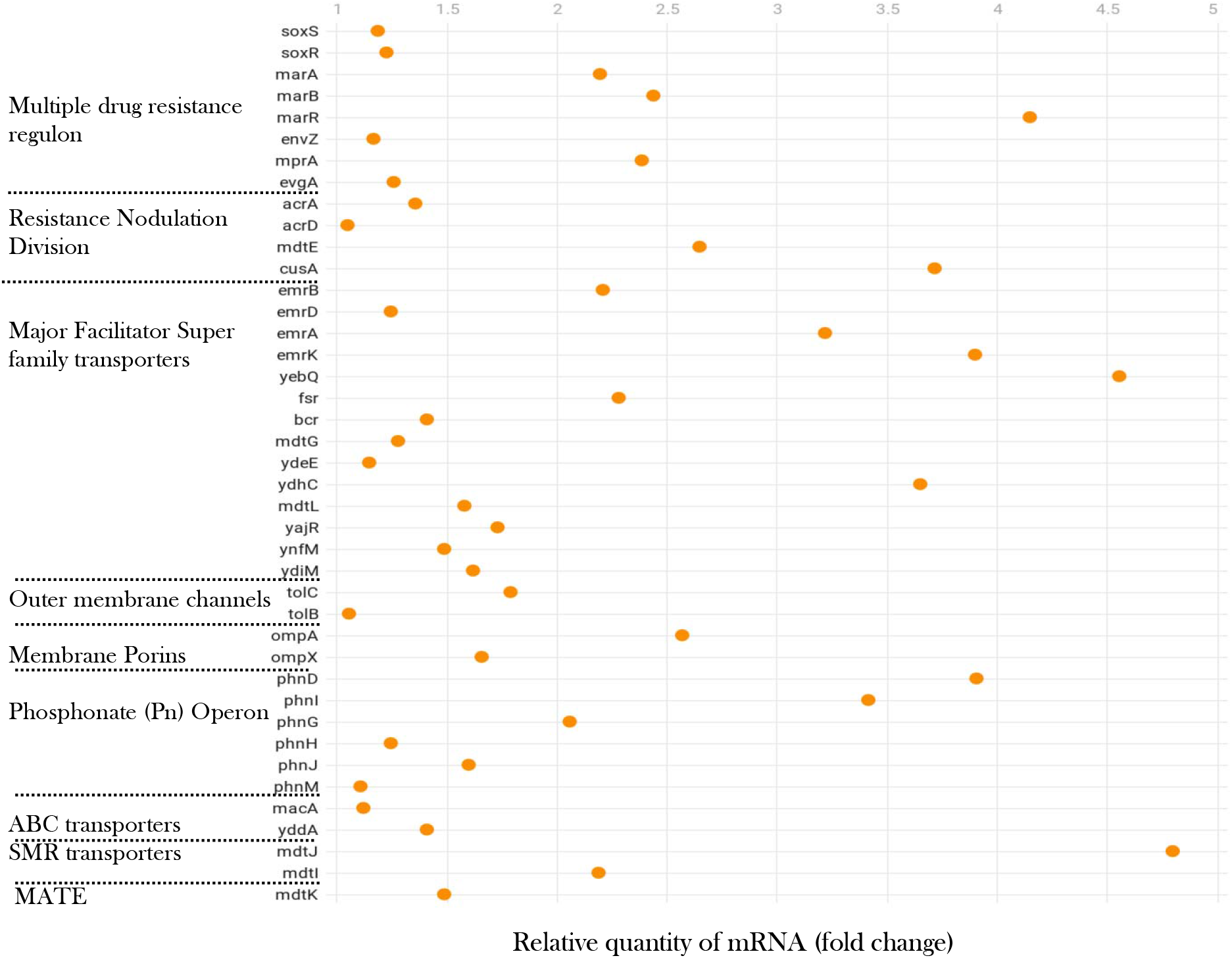
Complete list of differentially expressed transporters on CP exposure.

**Figure 5.6:**
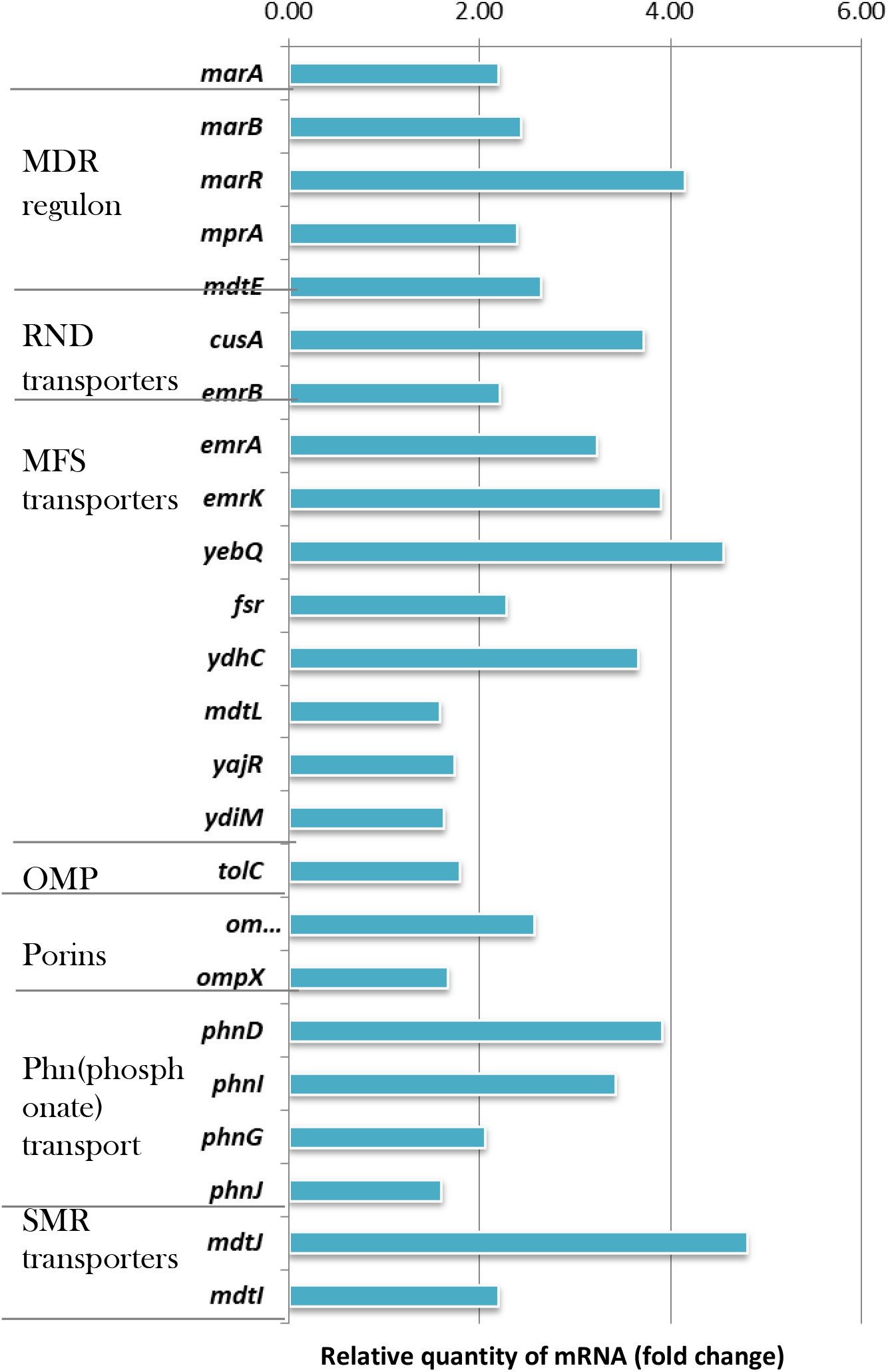
The transporters significantly expressed in the presence of chlorpyrifos in *E. coli*

**Figure 5.6:**
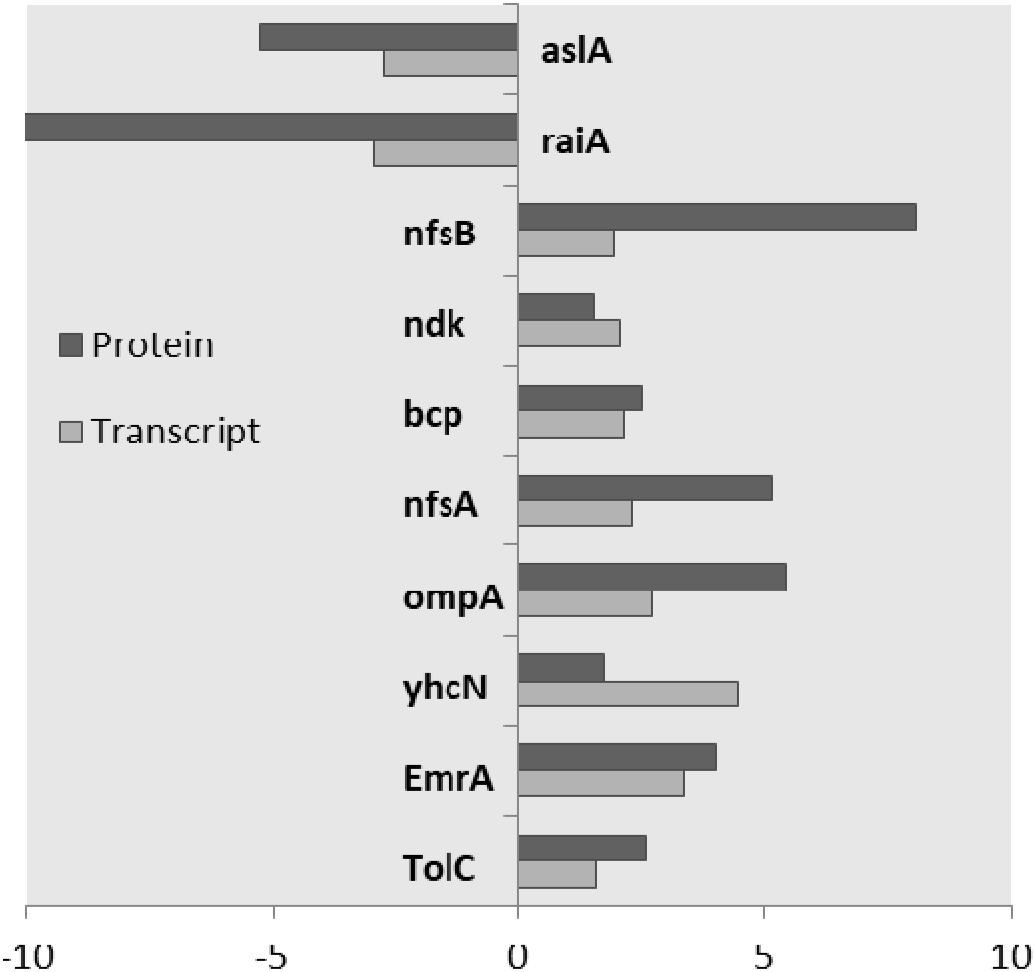
The most significantly expressed genes/proteins at both protein and transcript level on exposure to CP.

### 3.4. Global comparison of differential gene expression and protein expression on exposure to chlorpyrifos

The comparison of RNA-seq transcript expression profile with the differential proteome profile of *E. coli* on exposure to 500 mg/L of CP yielded 177 proteins that were differentially expressed at both the transcript and protein level. Of these, 72 showed significant differences in expression in either transcripts or proteins and were clustered into four categories. Of these, 24 showed significant expression in both the transcript and protein (1.5 >fold < −1.5), six showed significant expression in transcript but reduced expression in protein, while 29 displayed significant expression in protein but reduced expression in transcript and 11 revealed lower expression in both protein and transcript. The most significant differentially expressed entries in both transcript and RNA are given in Figure 5.7. Combined transcriptomic and proteomic approaches can provide clues on the adaptive mechanisms that come into play when the bacteria get exposed to an adverse environmental condition.

Though a direct correlation between the expressed RNA and protein could be expected, it needn’t always be the case (Vogel and Marcotte, 2012). The transcriptome may undergo rapid remodelling during a sudden spurt in growth, while it appears to be relatively static in the stationary phase. A higher dynamics of proteome compared to transcriptome was observed in *Rhodobacter sphaeroides* during the growth, and the organism responded to adverse environmental conditions mostly at the protein level. However, during the sudden spurts in the growth, the first response was a rapid remodelling of the transcriptome, followed by delayed changes in the proteome (Bathke et al., 2019). Low correlation between the transcriptome and proteome points to the role of post transcriptional modifications, changes in expressed proteome mediated through sRNAs. While a higher correlation between the proteome and transcriptome under stress were also shown (Berghoff et al., 2013), it could be observed that the response of bacteria to xenobiotic/environmental stress involves complex regulation at both transcriptional and post-transcriptional levels. This includes dynamic changes in the transcriptome and proteome ensuring survival under hostile environmental conditions.

Gram negative bacteria enjoy a wide collection of efflux pumps in its arsenal capable of transporting structurally diverse molecules in and out of bacterial cell. The most studied molecules are antibiotics and the efflux mechanisms helps to lower their intracellular concentration or the concentration of any other potentially toxic/harmful xenobiotic allowing the bacteria to survive under higher xenobiotic concentrations. Over-expression of efflux pumps has been regarded as an intrinsic defence mechanism in Gram-negative bacteria that provides them an increased antibiotic resistance. The efflux pumps capable of transporting more than one structural class of antimicrobial drugs/antibiotics are called Multi Resistance Drug Resistance (MDR) transporters and they play a major role in antibiotic resistance (Blair et al., 2014).

The genetic response of bacteria to organophosphate pesticides and the mechanism of their bactericidal action are poorly understood. One major efflux system of interest for CP tolerance is EmrAB-TolC. The emr locus, consisting of the open reading frames emrA (membrane fusion protein) and emrB (inner membrane component), is a multidrug resistant domain, capable of conferring resistance to uncouplers such as carbonylcyanide m-chlorophenylhydrazone (CCCP), tetrachlorosalicylanilide (TCS) and antibiotics like nalidixic acid. Disruption of *emr B* significantly decreased resistance to uncouplers (Lomovskaya and Lewis, 1992). They concluded that the *emr* locus conferred resistance to *E. coli* cells from the uncouplers of oxidative phosphorylation and highly hydrophobic compounds. Since CP is a hydrophobic OP pesticide and emr locus is known to confer resistance only against compounds with high hydrophobicity together with the fact that *E. coli* showed increased expression of EmrA at both transcript and protein level on exposure to CP, it can be postulated that EmrAB-TolC pump plays a predominant role in pesticide tolerance of Gram negative bacteria. Extensively studied and clinically significant pumps such as AcrAB-TolC (Du et al., 2014; Fernandez-Recio et al., 2004) belong to RND family, while little is known about the functionally similar Major Facilitator Superfamily (MFS) pump Emr AB-TolC even after being known since decades. The two are constitutively expressed pumps with overlapping substrate ranges and possibly have multiple substrate binding sites, allowing simultaneous handling of multiple drug substrates (Elkins and Mullis, 2007). While AcrAB-TolC is a trimer, EmrAB-TolC possibly exists as a dimer *in vitro* (Tanabe et al., 2009). Since the efficiency of AcrAB-TolC system is suggested to be lower in the transport of chemicals from the cytoplasm; other transporters are expected to play a major role in the case of drugs with intracellular targets (Schuldiner, 2018; Shuster et al., 2016; Tal and Schuldiner, 2009). Multi drug transporters are essential in acquiring high-level resistance against drugs, as seen in case of triple knockouts of MdfA, MdtM and EmrE failing to resist higher concentrations of quinolones, while acrB knockout did resist these antibiotics, hinting that acrB may sometimes be dispensable (Schuldiner, 2018). On exposure to CP, *E. coli* did not show a significant expression of acrB which might suggest its incapability to remove the pesticide from the cell.

The emrR acts as the negative regulator of emr locus comprising of emrRAB operon. An additional promoter for emrB is located in the structural part of emrA and its expression increased in the stationary phase. Since the operon is capable of handling structurally unrelated drugs/compounds, their intrinsic function was suggested to be the efflux of these chemicals in *E. coli* (Lomovskaya and Lewis, 1995). Proper regulation of emrR-emrAB operon is essential for the survival of Agrobacterium *tumefaciens* on exposure to toxins (Khemthong et al., 2019). Hence, it can be concluded that emr locus essentially protects and responsible for the high tolerance of *E. coli* to chlorpyrifos.

## 4. Conclusions

Tolerance to pesticides is a resultant of a combination of varying mechanisms, the major one being efflux. The inhibition of efflux pumps by a broad spectrum efflux inhibitor PAβN lead to the accumulation of chlorpyrifos in *E. coli* cells and correspondingly decreasing its tolerance to chlorpyrifos. Two main classes of transporters were observed to be over-expressed in the gene expression analysis, mainly the single component transporters MdtJ and Mdt I belonging the SMR family, and MdtK belonging to the MATE family, both of which move compounds from the cytoplasm to periplasm. Multi-component transporters, mainly the TolC-dependent transporters capable of transporting chemicals across both the membranes (Emr AB, MdtE and EmrK) were the other major class which was over expressed. These pumps work in coordination to accommodate wide range of xenobiotics and minimise the leakage of hydrophobic substances back into the cell. The major efflux pump that plays a significant role in pesticide tolerance is the Major Facilitator Superfamily pump, Emr AB-TolC. The pump is over-expressed/up-regulated at both the gene and protein level in the presence of CP. The impact of PAβN on the susceptibility of bacteria to non-antibiotics such as the organophosphate pesticide chlorpyrifos suggests that CP is a substrate of efflux pumps in Gram negative rods, i.e., *E coli*. Significant downregulation of multi drug resistance (MDR) efflux pumps in *Pseudomonas nitroreducens* AR-3, a bacterium capable of chlorpyrifos degradation was also observed as part of the current study (described in chapter 6). This implied that upregulation of efflux pumps enhanced the CP tolerance in no/low degrading Gram negatives such as *E coli,* while in degraders it was down-regulated.

